# Residual ellipticity in waveplate-compensated polarization-resolved SHG microscopy may arise from femtosecond laser spectral bandwidth

**DOI:** 10.64898/2026.02.24.707711

**Authors:** David Nguyen, Jeffrey P. Wilde, Virginie Uhlmann, Daniel James Smith, Justine Kusch-Wieser, Valeria Zanrè, Jakob Schwiedrzik, Gábor Csúcs

**Affiliations:** Scientific Center for Optical and Electron Microscopy (ScopeM), ETH Zürich, Switzerland; Department of Electrical Engineering, Stanford University, USA; Department of Molecular Life Sciences, University of Zürich, Switzerland; Laboratory for High Performance Ceramics, Empa Dübendorf, Switzerland

## Abstract

Polarization-resolved second harmonic generation microscopy provides structural information about non-centrosymmetric biological samples such as collagen. It involves illuminating the sample with a focused laser beam having a variable linear polarization angle and recording the second harmonic signal as a function of this angle. However, accurate linear polarization control is challenging due to ellipticity introduced by reflections from mirrors and dichroic mirrors in the optical path. Waveplate-based compensation has emerged as the standard approach to address these distortions, but its effectiveness for quantitative measurements remains incompletely characterized. Here, we attempt to fill this gap by implementing an established automated waveplate compensation method based on a rotating half-waveplate in combination with a compensating quarter-waveplate. This was done on a commercial Leica TCS SP8 MP multiphoton microscope, making various hardware improvements and carefully documenting important experimental details. Despite significant effort, we consistently observed substantial unwanted residual polarization ellipticity, with amplitudes up to 0.25, persisting under optimal waveplate configurations. Our simulation analysis provides evidence that this limitation may arise from wavelength-dependent dichroic mirror birefringence combined with the broad spectral bandwidth (10nm to 20nm full width at half maximum) of femtosecond laser pulses. While the approach investigated here can compensate a single wavelength, different spectral components within the pulse experience different phase retardations from wavelength-dependent optical elements, potentially resulting in residual ellipticity that cannot be eliminated. Our simulations qualitatively reproduced key features of the experimental observations. These findings have important implications for quantitative polarization-resolved second harmonic generation microscopy and suggest that alternative approaches, including specimen rotation or picosecond laser sources with narrower bandwidth, should be investigated for applications requiring precise polarization control. To facilitate community investigation of these effects, we provide open-source analysis code and simulation files.

## Introduction

Second harmonic generation (SHG) is a nonlinear optical process in which two identical photons combine to form a single photon at exactly twice the incident frequency [1]. An essential requirement for SHG is that the medium is non-centrosymmetric [2]. This symmetry requirement has made SHG particularly valuable for biological imaging, where non-centrosymmetric structures such as collagen can be studied without labels [3, 4]. The polarization dependence of SHG provides structural information about fiber orientation and molecular organization [5]. Polarization-resolved measurements are performed by illuminating the sample with linear polarization and varying the relative angle between the fundamental beam polarization and the specimen orientation while recording SHG intensity [6]. However, achieving accurate linear polarization control is challenging due to ellipticity introduced by reflections from mirrors and dichroic mirrors, and polarization distortions from high numerical aperture (NA) objectives [7]. In commercial multiphoton microscope systems, where the optical layout is largely fixed and polarization filtering immediately before the objective is generally not possible, polarization distortions must be actively managed. While the use of lower NA objectives minimizes these distortions, residual ellipticity from optical elements in the beam path must still be addressed. Two approaches exist: compensating with waveplates to enable variable incident polarization [8], or performing measurements at a fixed incident polarization state (chosen to introduce minimal ellipticity) while rotating the specimen to vary the relative orientation [9]. The waveplate compensation approach uses a linearly polarized laser beam passed through a quarter-waveplate (QWP) followed by a half-waveplate (HWP). When a principal axis of the QWP is aligned with the incident polarization, the beam remains linearly polarized and the HWP can be rotated to control the polarization angle on the sample. The QWP can then be adjusted to compensate for ellipticity introduced by optical elements between the waveplates and the specimen, with the goal of maintaining optimal linear polarization across the full range of HWP rotation angles. Specimen rotation, however, is complicated by alignment challenges, as the rotation axis is typically not coaxial with the imaging field, making waveplate compensation the more commonly adopted approach.

Building on the automated waveplate compensation method for commercial microscopes developed by Romijn et al. [10], we implemented their approach on a Leica TCS SP8 MP—the same instrument type tested in their work. We improved system characterization by implementing a fast motorized polarization analyzer with a photodiode detector, enabling high-resolution ellipticity maps with double the waveplate angular sampling resolution and significantly reduced acquisition time. Our systematic measurements revealed that compensation primarily symmetrizes the angle-dependent ellipticity, but substantial residual ellipticity persists under optimal configurations. We attribute this effect to a combination of wavelength-dependent dichroic retardance [11] with the broad spectral bandwidth of femtosecond lasers, highlighting a fundamental limitation of the waveplate compensation approach. In this work, we present our hardware and software implementation, systematic ellipticity measurements at the entrance port and sample plane, and computational simulations that support our hypothesis regarding the physical mechanisms underlying these limitations. These findings have important implications for quantitative polarization-resolved SHG microscopy.

## Materials and methods

The polarization rotation and compensation module was implemented following Romijn et al. [10] and consists of a QWP followed by a HWP in the fundamental beam path before the microscope (Fig 1). The fundamental wavelength was 880nm, generated by a Ti:Sapphire femtosecond laser (Mai Tai, Spectra-Physics). This waveplate compensation approach was originally developed by Chou et al. [8] using a HWP-QWP configuration. The waveplate ordering is critical as it determines the mathematical formulation of the compensation curves; the QWP-HWP order offers the practical advantage that the QWP can be fixed while HWP rotation controls the polarization angle. Both waveplates were mounted in motorized rotation stages (Table 1). Implemented in this way, the compensation module requires approximately 70mm of optical path length, plus additional space if enclosed for laser safety. We positioned it after beam conditioning optics and immediately before the microscope entrance port (see S1 Fig). Due to variability in beam delivery between commercial systems, we do not provide detailed mounting specifications. The waveplates were positioned approximately perpendicular to the beam axis and centered in the beam path.

**Table 1.** Compensation module main components.

| Component | Quantity | Model | Vendor |
| --- | --- | --- | --- |
| Achromatic quarter-waveplate | 1 | AQWP10M-980 | Thorlabs |
| Achromatic half-waveplate | 1 | AHWP10M-980 | Thorlabs |
| Motorized rotation stage with controller | 2 | KPRM1E/M | Thorlabs |

**Fig 1.**
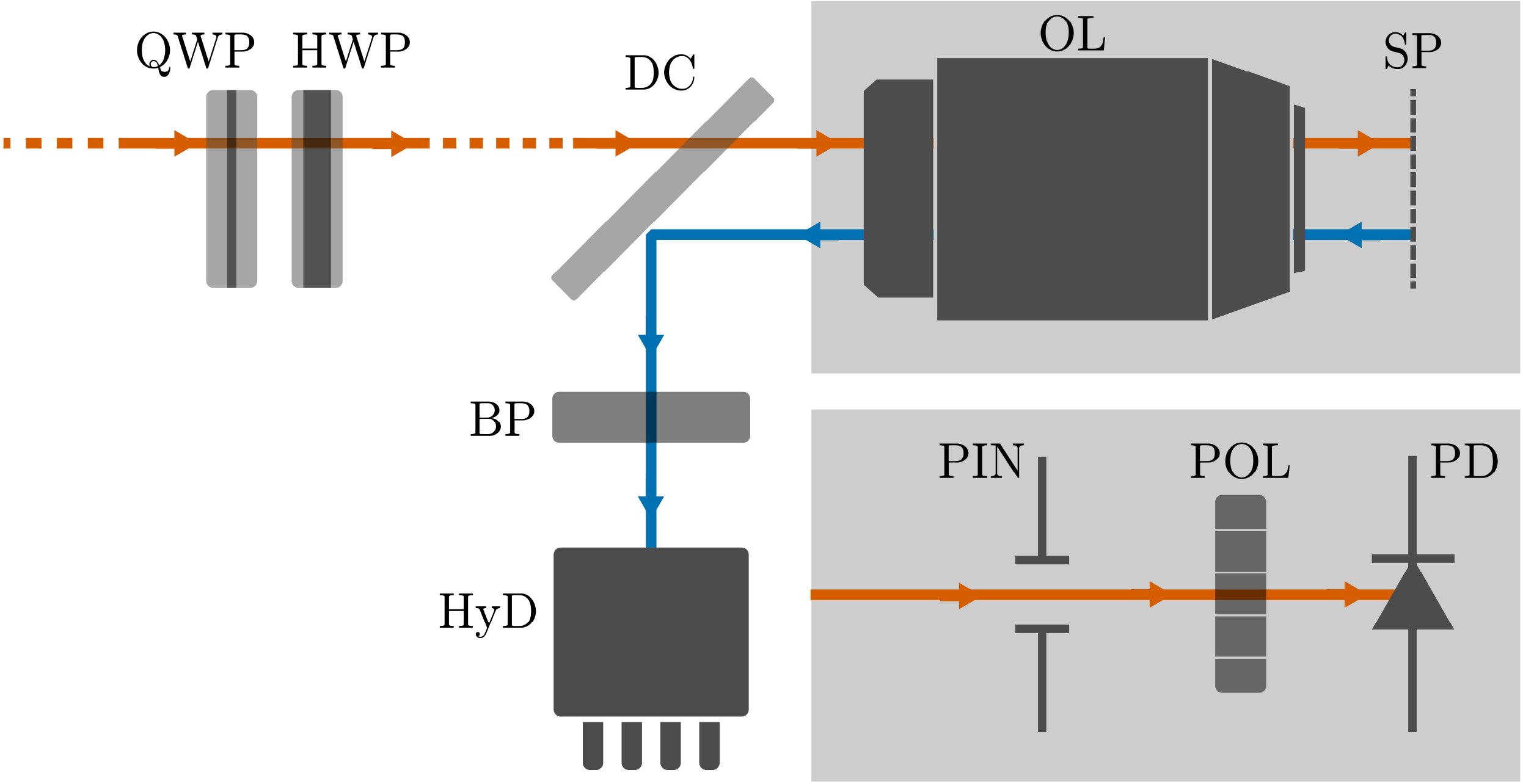
Optical layout. Fundamental beam from the laser tuned at 880nm (orange) passes through the beam conditioning optics (not shown, indicated by dashed line) before entering the compensation module made of a quarter-waveplate (QWP) and half-waveplate (HWP). The beam then passes through the microscope optics (dashed section) to the objective lens (OL) and is focused at the sample plane (SP). In imaging mode, back-scattered SHG signal (blue) is reflected by the dichroic mirror (DC) and detected in the non-descanned path through a bandpass emission filter (BP, FF01-445/20, Semrock) by a hybrid detector (HyD) from Leica. During calibration, the OL is removed and a polarization analyzer consisting of a pinhole (PIN), polarizer (POL), and photodiode (PD) is inserted.

The polarization analyzer is also based on the method proposed by Romijn et al. [10], although with substantially improved sampling rate and acquisition speed. Our system comprises four key components (Table 2): a motorized rotation stage capable of 5 Hz (1800 ^◦^*/*s) rotation speed, a polarizer mounted on the rotation stage, a photodiode for measuring transmitted power, and a 16-bit data acquisition (DAQ) device selected for increased dynamic range compared to lower bit-depth alternatives, which maximizes ellipticity measurement sensitivity. The mechanical assembly is illustrated in S2 Fig and computer-aided design (CAD) files are available in S1 File (SolidWorks Premium 2022 SP5.0, Dassault Systèmes).

**Table 2.** Polarization analyzer main components.

| Component | Model | Vendor |
| --- | --- | --- |
| Pinhole | P4000K1 | Thorlabs |
| Polarizer | LPVIS050-MP2 | Thorlabs |
| Motorized rotation stage | DDR25/M and KBD101 | Thorlabs |
| Photodiode | SM1PD1A | Thorlabs |
| Bias module | PBM42 | Thorlabs |
| Variable terminator | VT2 | Thorlabs |
| Data acquisition device | USB-6002 | NI |

The rotation stage completes individual turns in approximately 350 ms, including acceleration and deceleration phases. The stage controller outputs digital triggers at every degree from 0^◦^ to 359^◦^. Since the DAQ does not support retriggerable tasks (hardware-triggered finite acquisition that resets on each trigger), data are acquired continuously from both the photodiode and the trigger signal simultaneously. Acquisition runs for 360 ms at 25 000 samples*/*s per channel, yielding 9000 samples per channel. This rate is limited by the DAQ’s maximum aggregate sampling rate of 50 000 samples*/*s shared between channels. Post-processing identifies trigger positions in the recorded trigger channel to reconstruct the photodiode signal versus polarizer rotation angle. To reduce measurement variation, the sample at each trigger position is averaged with its two subsequent samples (angular displacement calculated in S1 Appendix). The complete electronics and wiring connections are shown in S3 Fig.

The DAQ analog input exhibits an offset that stabilizes after a 20 min warm-up period. This offset is recorded by averaging the channel readout for 6 s in complete darkness and typically ranges from − 2mV to − 3mV in our measurements. Both raw analog data and offset values are saved; background subtraction is performed during post-processing alongside trigger-based resampling. Operating parameters require careful consideration: photodiode linearity is verified by confirming proportional response to laser power variations, and the load resistance is maintained below 5 kΩ to ensure adequate bandwidth for the 25 ksamples s^−1^ acquisition rate. Detailed bandwidth calculations are provided in S2 Appendix. All measurements in this work employed a 500Ω load with 10V bias voltage, constrained by DAQ analog output specifications. Validation against a power meter demonstrated agreement for ellipticity values above approximately 0.074 ellipticity (see S4 Fig).

For comparison, a single ellipticity measurement in Romijn et al. [10] required approximately 17 s. While single measurement duration is not documented in this previous work, an analysis of the published code reveals 6 samples at 30^◦^ intervals (line 116 of `polarization_map.m`) over 0^◦^ to 150^◦^ (line 296 of `polarization_map.m`), with 1 s required for each power measurement and 25 ^◦^/s maximum rotation speed. Assuming a 25 ^◦^/s^2^ acceleration with a trapezoidal velocity profile yields an approximately 2.2 s travel time between samples. In contrast, our polarization analyzer completes a measurement in 360 ms, which constitutes a substantial improvement although the overall calibration time remains limited by the waveplate rotation stages (see Results and discussion).

### Software implementation

We provide the complete Python codebase for experimental data acquisition and analysis in S2 File (https://github.com/Omnistic/residual_ellipticity_in_pshg_hardware_and_analysis). The core computational routine implements the intensity fit for single polarization measurements following Eq 7 from Romijn et al. [10]. We use SciPy’s `curve_fit` function [12] with several modifications to improve numerical stability. The photodiode signal is scaled by a factor of 10 000 to avoid small numerical values from the DAQ (resolution is 0.305mV). To enforce the physical constraint *E*_*max*_ ≥ *E*_*min*_, we reparameterize using an intermediate variable *k* where *E*_*max*_ = *E*_*min*_ + *k*^2^, fitting the parameters (*α*_*max*_, *E*_*min*_, *k*) without requiring initial guesses. The bounds are set to (0,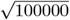) for (*E*_*min*_, *k*) based on the 10 V maximum analog input range, and (0, *π*) for *α*_*max*_ in radians.

For the ellipticity map fitting (Eq 1-3 in Romijn et al. [10]), we apply the same photodiode signal scaling factor. The bound for *I*_0_ = *E*_0_^2^ is set to (0, 100000). Note that *I*_0_ is missing from Eq 3 in Romijn et al. [10] but appears as *E*_0_ in their Eq 1 and as *I*_0_ in their code implementation (line 772 of `polarization_map.m`). The reflection ratio parameter *γ* only has a positivity constraint using the bounds (0, inf) and all the angles are constrained within (− *π, π*) in radians. A generic initial guess is provided as 10 000 for *I*_0_ (actual photodiode signal voltage on the order of 1V), 1.0 for *γ*, and 0^◦^ for all the angles. The optimal compensation curves that minimize ellipticity (Eq 4 in Romijn et al. [10]) are determined using SciPy’s `root` function. To ensure convergence to the correct solutions, the user supplies two initial QWP angles that minimize ellipticity on average across all HWP angles, identified by visual inspection of the ellipticity map.

## Results and discussion

To establish baseline performance, we first measured ellipticity as a function of relative polarization angle (*α*_*max*_) using only the HWP, without the QWP. Measurements were taken at both the entrance port (right after the HWP) and the sample plane (Fig 2A). At the entrance port, the ellipticity remains below 0.05. There is variation of the ellipticity with polarization angle, but as it falls within the laser polarization specification (given as an extinction ratio 500:1, corresponding to an ellipticity of 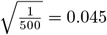) and at the edge of our detection sensitivity, we consider the polarization to be effectively linear at this point. By comparison, at the sample plane, the ellipticity shows periodic variation with two peaks and two valleys, ranging from 0.06 to 0.25.

**Fig 2.**
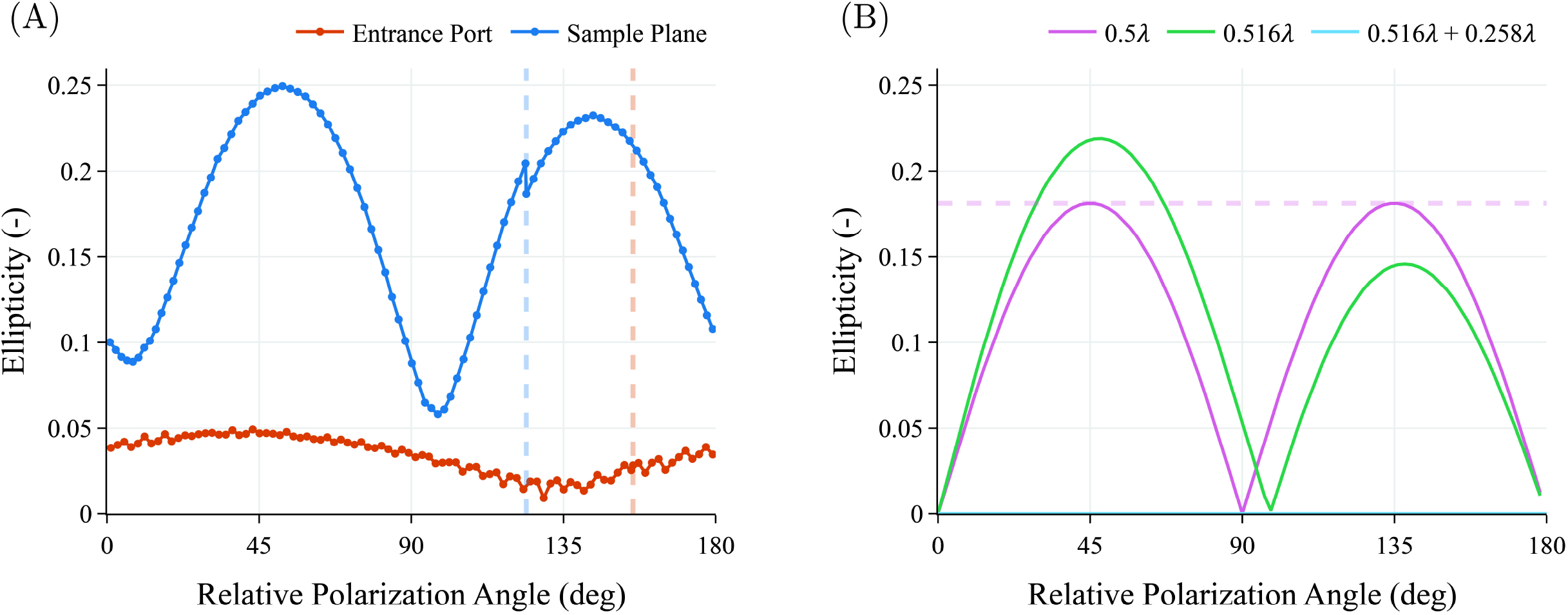
Preliminary ellipticity measurements and simulation. (A) Measured ellipticity as a function of relative polarization angle (*α*_*max*_) using only the HWP, without the QWP. Measurements were taken at the entrance port (right after the HWP, orange) and at the sample plane (blue). At the entrance port, ellipticity remains below 0.05, within laser specification. At the sample plane, ellipticity shows periodic variation with two peaks and two valleys, ranging from 0.06 to 0.25. Vertical dashed lines indicate the HWP motor zero position for each measurement. (B) Simulated ellipticity using a Jones matrix model in OpticStudio 2025 R1 (Ansys Zemax). The optical path consists of the QWP and HWP (modeled as linear retarders), followed by five silver-protected mirrors at 45^◦^ based on Chou et al. [8], and a dichroic mirror modeled as a linear retarder with 12.1^◦^ retardance. For the HWP-only simulations, the QWP retardance is set to zero. Simulation with ideal HWP (retardance = 0.5λ, pink curve) produces symmetric peaks and valleys, while simulation with manufacturer-specified HWP (retardance = 0.516λ, green curve) reproduces the asymmetry observed in the sample plane data (panel A), where peaks reach different heights and valleys reach different minima. A pink horizontal dashed line marks the first peak maximum to highlight this asymmetry. Adding the QWP with its manufacturer-specified retardance to the real HWP (light blue curve) demonstrates that ellipticity can be fully compensated despite non-ideal waveplate retardances.

To understand the ellipticity variation pattern at the sample plane, we consider the differential reflection properties of metallic mirrors at non-normal incidence. In our optical system, mirrors are generally positioned at 45^◦^ to the beam path. At each reflection, the beam’s polarization can be decomposed into *s*-polarized (perpendicular to the plane of incidence) and *p*-polarized (parallel to the plane of incidence) components, which experience different phase shifts upon reflection from the metallic surface (see S5 Fig). When the beam polarization is oriented as purely *s* or purely *p*, no ellipticity is introduced, giving rise to the two valleys in the pattern separated by 90^◦^ (*s* and *p* polarizations are orthogonal). The maximum ellipticity (peaks) occurs at intermediate angles, specifically halfway between the *s* and *p* orientations. Rotating the HWP from 0^◦^ to 90^◦^ sweeps through these different polarization orientations (0^◦^ to 180^◦^), producing the periodic pattern. While this explains the basic two-peak, two-valley structure, additional optical components—including the dichroic mirror and potentially other elements in the commercial microscope system—add further complexity. To validate our understanding, we performed a Jones matrix [13] simulation based on the optical model from Chou et al. [8] (illustrated in their figure 1), which consists of a cascade of five mirrors followed by a dichroic mirror. We modeled the mirrors as silver-protected reflectors to match our actual system, and the dichroic mirror as a linear retarder with 12.1^◦^ retardance, the value determined by the Romijn et al. [10] calibration method applied to our system (see below). The simulation was implemented in OpticStudio 2025 R1 (Ansys Zemax), where the HWP angle was swept programmatically using the ZOS-API via the open-source ZOSPy Python wrapper [14]. At each HWP angle, we calculated the output polarization state at the sample plane. Linear retarders (including the waveplates and dichroic mirror) were modeled as Jones matrix surfaces with the form:

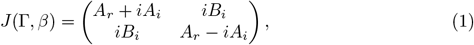

Where

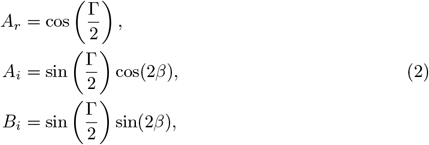

with Γ the retardance and *β* the fast axis angle of the retarder. To simplify parameter testing, we implemented these matrix elements using custom ZPL macro solves (one for each coefficient in Eq 2). The ZPL macro implementation is detailed in S3 Appendix, and OpticStudio files and simulation codes are provided in S3 File (https://github.com/Omnistic/residual_ellipticity_in_pshg_simulations).Figure 2B shows the simulated ellipticity pattern as a function of the relative polarization angle. With an ideal half-waveplate (retardance = 0.5λ, where λ is the wavelength), the simulation produces a periodic pattern ranging from 0.0 to 0.18 with symmetric peaks and valleys. However, using the manufacturer-specified retardance of 0.516λ at 880nm introduces asymmetry: the two peaks reach different heights and the second valley shifts from 90^◦^ to 98^◦^. This asymmetry matches the behavior observed in our experimental data. Importantly, the simulation demonstrates that even with non-ideal waveplate retardances, adding the QWP with a retardance of 0.258λ at 880nm enables complete compensation, reducing the ellipticity to zero across all angles (as shown by the light blue line that falls along the horizontal axis in Fig 2B).

We then experimentally applied the full compensation method of Romijn et al. [10], measuring ellipticity as a function of both QWP and HWP angles to create two-dimensional ellipticity maps at the entrance port and sample plane (Fig 3A). Our improved acquisition speed enabled maps with double the angular sampling resolution while reducing acquisition time from approximately 2 h to 30 min per map compared to the original implementation.

**Fig 3.**
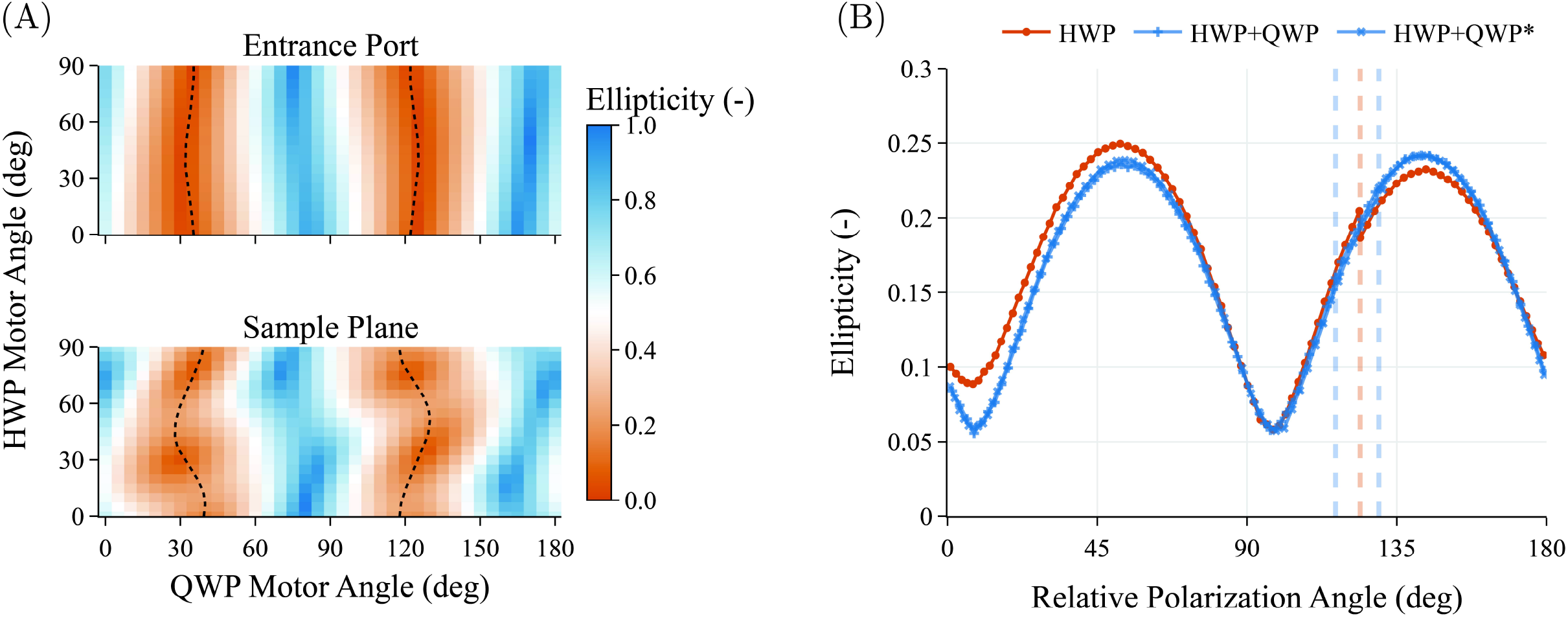
Two-dimensional ellipticity maps and compensation performance. (A) Ellipticity as a function of QWP and HWP angles at the entrance port (top) and sample plane (bottom), measured following Romijn et al. [10] with double the angular sampling resolution. Each map was acquired in approximately 30 min. At the entrance port, the map displays nearly vertical stripes indicating minimal ellipticity requiring small QWP corrections. At the sample plane, the pattern exhibits pronounced waviness with substantially larger QWP amplitude required for compensation. Dashed black lines indicate the optimal compensation curves (two solutions for achieving linear polarization at each HWP angle, calculated using Eq 4 from Romijn et al. [10]). Even along these optimal curves, ellipticity varies substantially, reaching values up to 0.25 in certain regions. (B) Ellipticity measured at the sample plane along the two optimal compensation curves (blue, extracted from the maps and resampled at 1^◦^ intervals) compared to the HWP-only baseline (orange). The first compensation solution (blue, + markers) and second solution (blue, × markers) overlap closely. While compensation produces more symmetric curves compared to the HWP-only baseline, substantial residual ellipticity persists as a function of HWP angle. The zero positions of the two compensation solutions are offset by approximately 13^◦^ (vertical dashed lines).

At the entrance port, the ellipticity map displays a pattern of nearly vertical stripes, indicating minimal ellipticity that requires only small QWP rotations for compensation. This is consistent with the single-HWP measurements (Fig 2A), confirming that the beam remains nearly linearly polarized at this location. In contrast, the sample plane map exhibits substantial distortion: the pattern develops pronounced horizontal waviness, and the amplitude of QWP rotation required for compensation increases significantly. This reflects the substantial ellipticity introduced by the optical path between the entrance port and sample plane.

Fitting the ellipticity maps to the model of Romijn et al. (Eq 1-3) provides estimates of key system parameters, though as discussed below, the model does not fully capture all sources of residual ellipticity. The fitted system retardance δ (representing cumulative birefringence from all optical elements after the compensation module) increases from 3.4^◦^ at the entrance port to 12.1^◦^ at the sample plane, confirming substantial birefringence accumulation through the microscope optics. As expected, the fitted QWP orientation angle *ϕ*_0_ remains nearly identical between measurement locations (33.7^◦^ and 33.9^◦^). The fitted HWP angle *θ*_0_ varies between the two locations (18.1^◦^ to 25.6^◦^). The fitted polarization analyzer angle *α*_0_ also differs between locations (11.3^◦^ vs. − 6.0^◦^), as expected since the polarization analyzer had to be mounted at different orientations at each location due to space constraints. Complete fitting parameters are provided in S1 Table.

The optimal compensation curves (dashed black lines), representing the two solutions for achieving linear polarization at each HWP angle, reveal a critical limitation: even where QWP and HWP angles are optimally adjusted, ellipticity varies substantially along these curves, reaching values up to approximately 0.25 at certain HWP angles while dropping near zero at others. To quantify this effect, we extracted these compensation curves from the ellipticity maps, resampled them at 1^◦^ intervals, and measured the resulting ellipticity at the sample plane (Fig 3B). While compensation produces more symmetric curves compared to the HWP-only case, substantial residual ellipticity persists and varies significantly as a function of HWP angle. The two compensation solutions overlap closely, with their zero positions offset by approximately 13^◦^ (indicated by vertical dashed lines). Both this shift and the residual ellipticity variation were already present in Romijn et al.’s data (their figure 6b and 6d), although not extensively discussed.

It should be noted that the acceptable level of ellipticity depends strongly on the specific application. Importantly, because ellipticity varies with polarization angle, it can modify the shape of polarization-dependent SHG intensity polar plots. Since shape parameters in these plots are commonly used to infer sample orientation and microstructural organization, such angle-dependent distortions can directly influence orientation-related analysis. For some quantitative polarization-resolved measurements, a residual ellipticity of 0.25 may be acceptable depending on the required measurement precision for subsequent analysis. However, this value is still substantial, particularly given that ellipticity as low as 0.06 can be achieved at certain relative polarization angles. Our key finding is that the waveplate compensation approach provided minimal improvement in our system: while ellipticity varied from 0.06 to 0.25 with HWP only, the residual ellipticity after full compensation remained comparable, simply with improved symmetry. This suggests that, for systems with relatively low retardance to compensate as in our case, the practical benefit of QWP compensation is limited.

We hypothesized that this residual ellipticity arises from the combination of wavelength-dependent dichroic mirror birefringence and the broad spectral bandwidth of femtosecond laser pulses. Dichroic mirror birefringence is well documented in the literature [11], including its strong wavelength dependence. Schön et al. measured dichroic mirror retardance using ellipsometry, demonstrating variations exceeding 50^◦^ over spectral ranges of 10nm to 20nm (their figure 6b).

This wavelength dependence becomes particularly relevant for polarization-resolved SHG microscopy because femtosecond laser sources possess relatively large spectral bandwidths. Our Spectra-Physics Mai Tai laser at 880nm has a specified bandwidth of 12.5nm (FWHM), comparable to the spectral range over which dichroic retardance can vary by tens of degrees. While a single retardance value can be compensated by appropriate QWP and HWP settings, each wavelength component within the pulse bandwidth experiences different retardance from the dichroic mirror. The waveplates, however, can only be optimized for a single wavelength or narrow spectral range. Consequently, wavelength components away from the optimization point accumulate different amounts of ellipticity, resulting in residual ellipticity that cannot be eliminated by any fixed waveplate configuration.

To explore this hypothesis, we extended our previous Jones matrix simulations to incorporate the polychromatic nature of femtosecond pulses. We modeled the laser spectral bandwidth as nine discrete wavelengths with weights following a Gaussian distribution (FWHM = 20nm, centered at 880nm; Fig 4A). While our laser is specified at 12.5nm FWHM, we used 20nm as a representative bandwidth for femtosecond pulses to explore the general effect of spectral bandwidth on residual ellipticity. The dichroic mirror retardance was modeled to vary linearly from − 12.9^◦^ to 37.1^◦^ across this bandwidth, centered at 12.1^◦^ at 880nm, spanning a total range of 50^◦^ consistent with the measurements of Schön et al. [11] (Fig 4B). For each QWP and HWP angle combination, we calculated the transmitted intensity as a function of polarization analyzer angle for all nine wavelengths independently, then summed these intensity curves incoherently to simulate the detector response. Ellipticity was then extracted from the summed intensity curve using the same fitting procedure as for experimental data (ellipticity extraction details in S4 Appendix). We provide the simulation code and OpticStudio files in S3 File.

**Fig 4.**
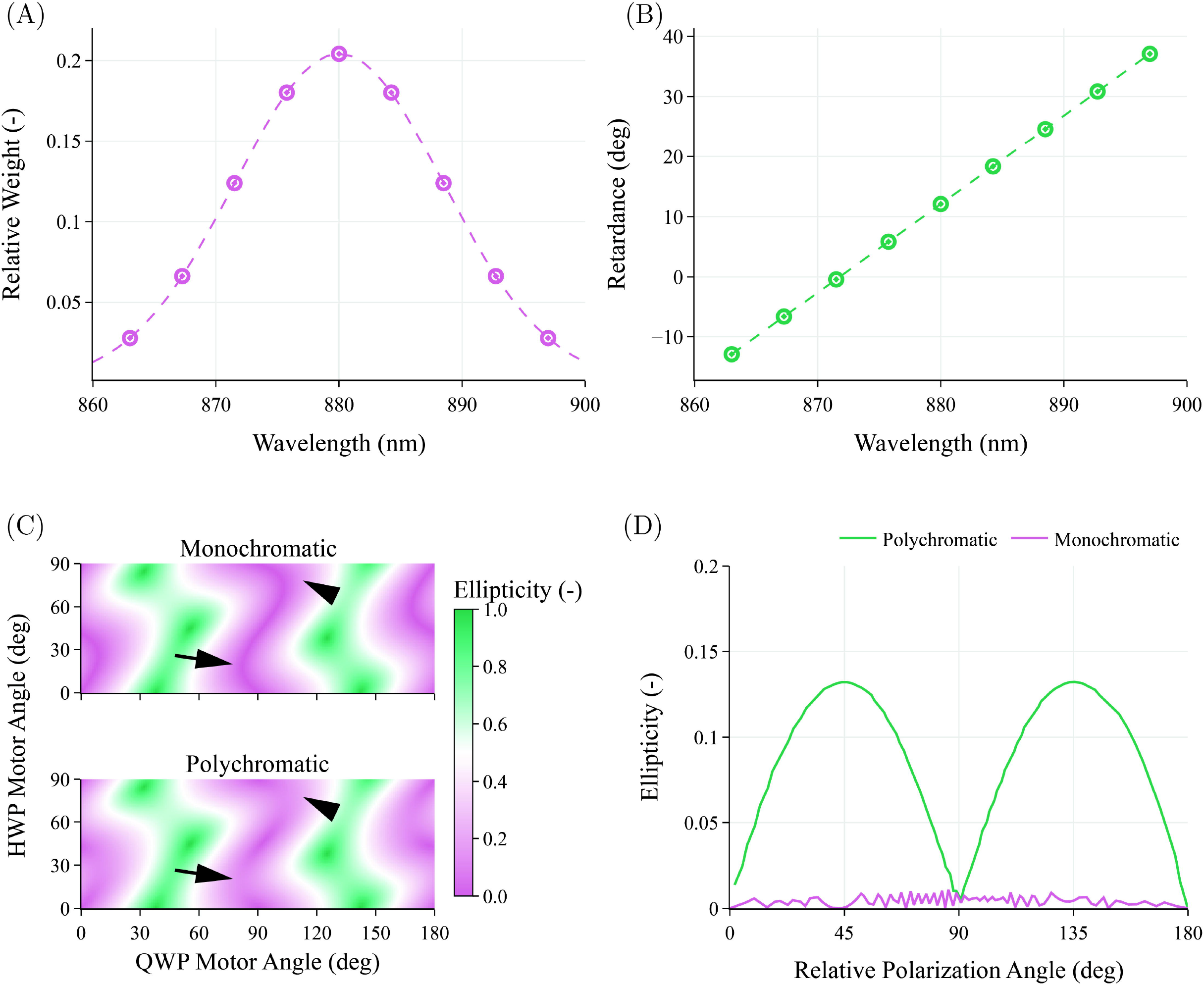
Polychromatic simulation reveals spectral origin of residual ellipticity after quarter-waveplate compensation. (A) Laser spectral bandwidth modeled as nine discrete wavelengths with weights following a Gaussian distribution (FWHM = 20nm, centered at 880nm). (B) Dichroic mirror retardance as a function of wavelength, modeled to vary linearly from − 12.9^◦^ to 37.1^◦^ across the laser bandwidth (total range of 50^◦^), centered at 12.1^◦^ at 880nm, consistent with variations in Schön et al. [11] (their figure 6b). (C) Simulated ellipticity maps for monochromatic illumination (880nm only, top) and polychromatic illumination (bottom). The optimal compensation curves are not displayed to better visualize the ellipticity modulation pattern. While monochromatic illumination achieves near-zero ellipticity along the compensation curves, polychromatic illumination exhibits two distinct peaks of residual ellipticity (marked by arrowhead and arrow). (D) Ellipticity extracted along the optimal compensation curve as a function of relative polarization angle for monochromatic (pink curve) and polychromatic (green curve) cases. The polychromatic simulation produces periodic residual ellipticity variation with amplitude of approximately 0.13, qualitatively reproducing the experimental observations in Fig 3B.

Figure 4C compares the resulting ellipticity maps for monochromatic (880nm only) and polychromatic illumination. The simulation reveals a striking effect: incorporating spectral bandwidth introduces periodic modulation of ellipticity along the optimal compensation curves. The curves are not displayed in the maps to better visualize this modulation pattern. While the monochromatic case achieves near-zero ellipticity along these curves, the polychromatic case exhibits two distinct peaks of residual ellipticity (marked by arrowhead and arrow in Fig 4C). This pattern is more clearly visible when plotting ellipticity along the compensation curve as a function of relative polarization angle (Fig 4D). The polychromatic simulation produces residual ellipticity variation with an amplitude of approximately 0.13, qualitatively reproducing the periodic variation observed in our experimental data (Fig 3B), though with lower amplitude.

While our simulation does not quantitatively match the experimental amplitude (which reaches approximately 0.25), it provides strong evidence that wavelength-dependent dichroic retardance combined with femtosecond pulse bandwidth represents a fundamental limitation of waveplate-based compensation. The discrepancy in amplitude could arise from several factors: our simplified linear retardance model, uncertainty in the actual dichroic retardance spectrum of our specific microscope, contributions from other wavelength-dependent optical elements, or the approximation of the spectral distribution. Direct measurement of our dichroic mirror retardance spectrum would be required for quantitative validation but is beyond the scope of this work.

## Conclusions

In this work, we have implemented and characterized the waveplate-based polarization compensation method previously proposed by Romijn et al. [10] on a commercial Leica TCS SP8 MP multiphoton microscope, and introduced modifications to achieve a substantially improved acquisition speed enabling high-resolution ellipticity mapping as a function of the QWP and HWP rotation angles. Our systematic measurements confirm that the compensation approach successfully symmetrizes angle-dependent ellipticity patterns. However, substantial residual ellipticity persists even under optimal waveplate configurations, varying periodically as a function of output polarization angle with amplitudes up to 0.25.

Through Jones matrix simulations incorporating polychromatic illumination, we provided evidence that this limitation arises from the combination of wavelength-dependent dichroic mirror birefringence and the broad spectral bandwidth (10nm to 20nm FWHM) characteristic of femtosecond laser pulses. While each wavelength component can theoretically be compensated, fixed waveplates can only optimize for a single wavelength, thus leaving other spectral components with residual ellipticity. This illustrates a key limitation of waveplate-based compensation for femtosecond laser systems rather than an implementation issue. It should be noted that our microscope exhibited relatively modest system retardance (12.1^◦^ at the sample plane). Systems with substantially higher birefringence may see more dramatic improvement from waveplate compensation (bringing ellipticity down from very high values) even though the residual ellipticity variation we observed would still persist. In our case, the initially low ellipticity meant that the compensation approach offered limited practical benefit.

These findings have important implications for quantitative polarization-resolved SHG microscopy. For applications requiring precise polarization control, we suggest two potential approaches for future investigation. First, specimen rotation: rather than attempting to compensate the beam polarization across all angles through the microscope optics, one could minimize ellipticity at the sample plane (with a QWP) for a single beam polarization orientation and instead rotate the specimen. While this approach introduces technical challenges in maintaining the field of view relative to the rotation axis, necessitating motorized x-y translation and rotation stages, it would circumvent the wavelength-dependent retardance problem entirely. Second, picosecond laser sources with significantly narrower spectral bandwidths could reduce the wavelength-dependent effects we have identified. While the longer pulse duration would reduce SHG generation efficiency, the improved polarization control may benefit quantitative applications where signal-to-noise is less limiting than polarization accuracy. Additionally, if feasible, achromatic dichroic mirrors designed for minimal retardance variation across the 10-20 nm bandwidth of femtosecond laser pulses at the operating wavelength could address the wavelength-dependent limitations identified here.

## Supporting information

**S1 Fig. Compensation module installation**. Photograph of the compensation module showing motorized rotation stages for the quarter-waveplate (QWP, orange arrow) and half-waveplate (HWP, blue arrow) positioned in the beam path before the microscope entrance port. The polarization analyzer is shown mounted immediately after the compensation module for entrance port measurements; the same analyzer was also positioned at the sample plane for sample plane measurements (not shown). Enclosures are open for the photograph; all components are enclosed during measurements for laser safety.

**S2 Fig. Mechanical assembly of the polarization analyzer**. CAD assembly of the polarization analyzer. Arrows indicate component positions with corresponding Thorlabs model numbers labeled. Component models were obtained from Thorlabs. The assembly shows the photodiode detector (SM1PD1A) mounted in the optical path; a power meter sensor could be interchanged using a quick mount (SM1QA) for validation measurements. Complete SolidWorks assembly files are provided in the supplementary materials archive. Electronics, wiring, and data acquisition connections are shown in S3 Fig.

**S3 Fig. Electronics and control system schematic**. Wiring diagram showing the photodiode detector (SM1PD1A) with bias module (PBM42) and variable terminator (VT2), data acquisition device (USB-6002), and motor control architecture. The photodiode operates with 10V bias and 500Ω load. The DAQ provides the DC bias voltage to the PBM42 via analog output AO0, limited to 10V but sufficient for this application. Motor controllers (KBD101 for analyzer rotation; 2× KDC101 for waveplates) connect via KEH3 hub for single USB interface. The KBD101 digital trigger output serves two purposes: it initiates the DAQ acquisition task (DI0) and is simultaneously recorded as an analog input (AI0) alongside the photodiode signal (AI1). This dual connection compensates for the DAQ’s lack of retriggerable task support, enabling precise synchronization of photodiode measurements with angular position (see Materials and methods).

**S4 Fig. Validation of photodiode against power meter**. Comparison of ellipticity measurements using the photodiode detector (SM1PD1A, orange) and a power meter (Thorlabs PM100D with S121C sensor, blue) across a range of ellipticity values. The S121C sensor is also photodiode-based with similar specifications to the SM1PD1A, enabling direct comparison. Both detectors feature large photosensitive areas, providing tolerance to alignment variations. To validate the photodiode across different ellipticity levels, a single QWP (without HWP) was positioned in the beam path at various fast axis angles relative to the beam polarization to deliberately introduce controlled amounts of ellipticity. Each panel shows boxplots of repeated measurements: 20 measurements per detector without the QWP (NO QWP), and 10 measurements per detector with the QWP at fast axis angles of 0^◦^, 5^◦^, 10^◦^, 15^◦^, 30^◦^, and 45^◦^. The y-axis of each panel is centered on the power meter median value (bolded) to facilitate visual comparison. Both detectors show excellent agreement at ellipticity values above approximately 0.074, with maximum differences below 0.003. Below this threshold, the photodiode systematically underestimates ellipticity compared to the power meter, likely due to the power meter’s dynamic range adjustment capability which enables more accurate detection at lower signal levels at the cost of measurement speed. The ellipticity range encountered in this work (0.06-0.25) lies predominantly above the 0.074 threshold, ensuring reliable measurements with the photodiode detector.

**S5 Fig. Linear polarization can become elliptical after metallic mirror reflection**. (A) Schematic showing a light ray reflecting from a mirror surface at 45^◦^ incidence. The beam’s linear polarization can be decomposed into *s*-polarized (orange arrows, perpendicular to the plane of incidence) and *p*-polarized (blue arrows, parallel to the plane of incidence) components. These components experience different phase shifts upon reflection, potentially introducing ellipticity in the polarization. (B) Phase shift versus angle of incidence for *s* and *p* polarization components reflecting from pure silver at 880nm (optical constants from https://refractiveindex.info, Shelf:MAIN, Book:Ag (Silver), Page:Jiang et al. 2016). At 45^◦^ incidence (highlighted), the phase difference is 13.7^◦^. This phase difference is what causes linear polarization to become elliptical. Calculation code available at https://github.com/Omnistic/fresnel_equations_with_metals. Actual optical simulations (Fig 2 and 4) use protected silver coatings from OpticStudio 2025 R1 (Ansys Zemax).

**S1 File. SolidWorks assembly files for polarization analyzer**. Archive containing the polarization analyzer CAD assembly shown in S2 Fig. The main files are `polarization_analyzer.sldasm` (assembly) and `polarization_analyzer.slddrw` (drawing). All component models were obtained from the Thorlabs website and are included via SolidWorks Pack and Go. Files were created with SolidWorks Premium 2022 SP5.0 (Dassault Systèmes). Available at https://doi.org/10.5281/zenodo.17736144.

**S2 File. Python code for data acquisition and analysis**. GitHub repository containing the complete experimental code base for polarization analyzer control, data acquisition, and ellipticity analysis. Includes implementations of fitting routines (Eq 7, Eq 1-3 from Romijn et al. [10]) with numerical stability improvements, polarization parameter calculations, and figure generation scripts. Available at https://github.com/Omnistic/residual_ellipticity_in_pshg_hardware_and_analysis.

**S3 File. OpticStudio simulation files and Python automation code**. GitHub repository containing OpticStudio 2025 R1 (Ansys Zemax) simulation files and Python code for programmatic control and analysis. Includes: (1) ZAR archive with complete OpticStudio lens files implementing Jones matrix simulations for monochromatic and polychromatic cases, including all ZPL macros for Jones matrix surface calculations, (2) Python scripts using ZOS-API and ZOSPy to automate component rotation and data extraction from OpticStudio, and (3) data processing and figure generation scripts for ellipticity analysis. The logic and implementation details of the ZPL macros used in OpticStudio are documented in S3 Appendix. Available at https://github.com/Omnistic/residual_ellipticity_in_pshg_simulations.

**S4 File. Experimental ellipticity measurement data**. Complete dataset of experimental measurements including two-dimensional ellipticity maps, baseline measurements, and validation data. Each measurement is stored as a NumPy compressed archive (.npz) with the following structure: `analog_data` contains raw analog input data (2 channels × 9000 samples) from the photodiode signal (channel 0) and motor trigger signal (channel 1) without offset subtraction; `measurement_data` contains processed photodiode intensity as a function of polarization analyzer angle (2 × 360 array); `calibration_mean` contains the dark current offset value used for background subtraction during post-processing; `calibration_std` contains the dark current standard deviation recorded as an indicator of measurement stability during DAQ warm-up. File naming conventions: timestamps with optional descriptions (YYYYMMDDTHHMMSSZ description.npz format) for individual measurements, HWP and QWP angles (HWP-XXX QWP-YYY.npz format) for full ellipticity maps, or HWP angle only (XXX.npz format) for HWP-only baseline measurements. Data can be processed using the analysis code in S2 File. Total size approximately 120MB, available at https://doi.org/10.5281/zenodo.18433941.

**S1 Appendix. Angular displacement during three-sample averaging**. This appendix calculates the angular displacement of the polarization analyzer rotation stage during the three-sample averaging window used in post-processing.

The data acquisition device samples at 25 000 samples*/*s, corresponding to a sampling interval of

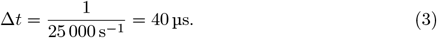

Post-processing averages each trigger position sample with its two subsequent samples, spanning a time window of

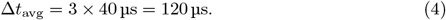

The rotation stage operates at a maximum speed of 1800 ^◦^*/*s. During the averaging window, the angular displacement corresponds to

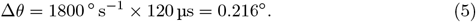

This angular displacement is small compared to the 1^◦^ trigger spacing and does not compromise measurement accuracy. The averaging improves signal-to-noise ratio while maintaining adequate angular resolution for ellipticity determination.

**S2 Appendix. Photodiode bandwidth calculation**. This appendix provides the calculation for the photodiode bandwidth to verify adequate frequency response for the polarization analyzer measurements.

The bandwidth of a photodiode detector is limited by the junction capacitance *C*_*j*_ and load resistance *R*_load_ according to:

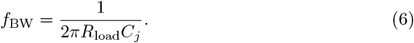

For the SM1PD1A photodiode operating at 10 V bias voltage, the manufacturer (Thorlabs) specifies a typical junction capacitance of *C*_*j*_ = 328 pF. With the load resistance of *R*_load_ = 500 Ω used in this work, the bandwidth is calculated as

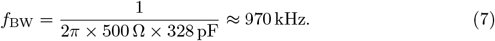

The data acquisition device samples at 25 000 samples*/*s, corresponding to a Nyquist frequency of 12.5 kHz. The photodiode bandwidth (*f*_BW_ = 970 kHz) well exceeds this requirement. At higher load resistances, such as *R*_load_ = 5 kΩ, the bandwidth would be approximately *f*_BW_ = 97 kHz, which still satisfies the Shannon-Nyquist criterion with comfortable margin. We recommend maintaining load resistance below 5 kΩ to ensure adequate bandwidth, as real-world signal integrity benefits from bandwidths substantially higher than the theoretical Shannon-Nyquist minimum.

**S3 Appendix. ZPL macro implementation for Jones matrix surfaces in OpticStudio**. This appendix details the implementation of Jones matrix surfaces in OpticStudio (Ansys Zemax) for modeling linear retarders such as waveplates and dichroic mirrors.

### Parameter reduction using physical constraints

OpticStudio’s Jones matrix surface type accepts eight parameters: the real and imaginary components of four complex coefficients (*A, B, C, D*). However, the physical form of a linear retarder given by Eq 1 shows that the matrix can be fully specified by only three independent values, as given in Eq 2. We implemented three ZPL macro solves to automatically calculate these coefficients from the physically meaningful parameters: retardance (Γ) and fast axis angle (*β*). These macros are embedded in the OpticStudio files and are automatically available when the ZAR archive (S3 File) is opened.

### ZPL macro solve implementation

The input parameters for each Jones matrix surface are specified using a comment-based system with dummy surfaces. Each Jones matrix surface is assigned a comment identifier (e.g., “QWP” for quarter-waveplate). Two dummy surfaces are placed immediately before the Jones matrix surface with comments “[IDENTIFIER] RETARDANCE” and “[IDENTIFIER] ANGLE” (e.g., “QWP RETARDANCE” and “QWP ANGLE”). The ZPL macro searches for these surfaces by parsing comments and reads the thickness values as the retardance (in degrees) and fast axis angle (in degrees), respectively. Since simulations trace a single ray coaxial with the optical axis, the additional optical path length from dummy surface thickness does not affect results, and these surfaces can be hidden in OpticStudio’s 3D Layout and Shaded Model views.

### Usage and benefits

This implementation enables direct manipulation of retardance and orientation in the OpticStudio interface without requiring ZOS-API or programmatic control. Users can rotate components or adjust retardance by simply modifying the thickness values of dummy surfaces, with all matrix elements updating automatically through the ZPL macro solves. This approach was used to implement the QWP, HWP, and dichroic mirror in all simulations presented in the main text. Complete ZPL macro code and example OpticStudio files are provided in S3 File.

### ZPL macro code

The three ZPL macros implement the coefficient calculations from Eq 2. Each macro follows the same structure: parse the comment to identify the surface, locate the dummy surfaces for retardance and angle, and compute the appropriate coefficient.

- Macro for *A*_*r*_ = cos(Γ*/*2):

~~~
CONS_PI = 2 * ACOS(0)
TO_RAD = CONS_PI / 180
current_surface_number = SOSO(0)
plate_name$ = $COMMENT(current_surface_number)
# Case-Insensitive
retardance_comment$ = plate_name$ + “_retardance”
retardance = THIC(SURC(retardance_comment$))
retardance = retardance * TO_RAD
SOLVE
RETURN COSI(retardance/2)
~~~

- Macro for *A*_*i*_ = sin(Γ*/*2) cos(2*β*):

~~~
CONS_PI = 2 * ACOS(0)
TO_RAD = CONS_PI / 180
current_surface_number = SOSO(0)
plate_name$ = $COMMENT(current_surface_number)
# Case-Insensitive
retardance_comment$ = plate_name$ + “_retardance”
angle_comment$ = plate_name$ + “_angle”
retardance = THIC(SURC(retardance_comment$))
angle = THIC(SURC(angle_comment$))
retardance = retardance * TO_RAD angle = angle * TO_RAD
SOLVE
RETURN SINE(retardance/2) * COSI(2*angle)
~~~

- Macro for *B*_*i*_ = sin(Γ*/*2) sin(2*β*):

~~~
CONS_PI = 2 * ACOS(0)
TO_RAD = CONS_PI / 180
current_surface_number = SOSO(0)
plate_name$ = $COMMENT(current_surface_number)
# Case-Insensitive
retardance_comment$ = plate_name$ + “_retardance”
angle_comment$ = plate_name$ + “_angle”
retardance = THIC(SURC(retardance_comment$))
angle = THIC(SURC(angle_comment$))
retardance = retardance * TO_RAD angle = angle * TO_RAD
SOLVE
RETURN SINE(retardance/2) * SINE(2*angle)
~~~

## S4 Appendix. Ellipticity extraction from OpticStudio simulations

This appendix describes how ellipticity and polarization angle are extracted from OpticStudio simulations for monochromatic and polychromatic cases.

For monochromatic simulations, ellipticity and polarization angle are calculated directly in OpticStudio using the CODA Merit function operand. The operand is configured with Data = 121, 122, and 123 to extract the major axis (*E*_*max*_), minor axis (*E*_*min*_), and angle (in degrees) of the polarization ellipse, respectively, as defined in the OpticStudio documentation. Ellipticity is computed as *E*_*min*_*/E*_*max*_, and the relative polarization angle (*α*_*max*_) is obtained directly from Data 123.

For polychromatic simulations, the CODA operand cannot be used directly because the polarization state depends on wavelength. A multi-configuration setup is used in OpticStudio with each configuration representing one wavelength, its associated Gaussian weight (Fig 4A), and the corresponding dichroic mirror retardance (Fig 4B). The transmitted intensity through a rotating linear polarizer as a function of polarization analyzer angle is calculated for each of the nine wavelengths independently using Eq 7 from Romijn et al. [10]. These intensity curves are then summed incoherently across all wavelengths to simulate the detector response to broadband illumination.

Ellipticity and polarization angle are extracted by fitting Eq 7 to the summed intensity curve using the same procedure as for experimental data (see Materials and methods).

**Table S1.** Fitted parameters from ellipticity maps. Parameters obtained by fitting the ellipticity maps (Fig 3A) to the model of Romijn et al. (Eq 1-3). Values are provided for measurements at both the entrance port (after the compensation module) and the sample plane. *I*_0_ represents the reference intensity, *γ* is the reflection ratio parameter, δ is the cumulative system retardance, and *θ*_0_, *ϕ*_0_, and *α*_0_ represent the angular offsets of the HWP fast axis, QWP fast axis, and polarization analyzer transmission axis, respectively, relative to their motor zero positions. The QWP orientation angle *ϕ*_0_ remains consistent between measurement locations (33.69^◦^ and 33.85^◦^), as expected since the QWP mounting was not changed. The HWP angle *θ*_0_ shows larger variation (18.07^◦^ to 25.61^◦^); this discrepancy may reflect the model’s incomplete capture of residual ellipticity sources as discussed in the main text. The polarization analyzer angle *α*_0_ differs between locations (11.28^◦^ vs. − 6.00^◦^) due to space constraints requiring different mounting orientations at each measurement position.

| Parameter | Entrance Port | Sample Plane |
| --- | --- | --- |
| $I_0$ | 2.01 | 1.96 |
| $\gamma$ | 0.99 | 1.07 |
| $\delta$ (°) | 3.40 | 12.10 |
| $\theta_0$ (°) | 18.07 | 25.61 |
| $\phi_0$ (°) | 33.69 | 33.85 |
| $\alpha_0$ (°) | 11.28 | −6.00 |

## Acknowledgments

The authors thank Elisabeth I. Romijn for helpful discussions regarding the implementation of the compensation method, as well as for clarifying the differences in waveplate ordering (QWP-HWP versus HWP-QWP) and the corresponding equations compared to the previous work of Chou et al [8].

We thank users `ZYOng` and `Kevin_Price` from the National Instruments Community Forums for helpful discussions regarding data acquisition with the USB-6002. `ZYOng` ’s suggestion to record the trigger signal as an analog input channel alongside the photodiode signal enabled high-speed synchronized acquisition without retriggerable task support (https://forums.ni.com/t5/Multifunction-DAQ/Non-uniform-retriggering-of-an-AI-task-with-USB-6002/td-p/4365407).

We thank the NiceGUI development team for helpful discussions regarding graphical user interface implementation (https://github.com/zauberzeug/nicegui/discussions/2985).

The authors used Claude 4.5 Sonnet (Anthropic) to assist with manuscript preparation, including structuring arguments, refining technical descriptions, and improving clarity. All scientific content, data analysis, interpretations, and conclusions represent the authors’ own work and ideas. All LLM-generated text was reviewed and edited by the authors for accuracy and appropriateness.

