## Supplementary figures and images for "Residual ellipticity in waveplate-compensated polarization-resolved SHG microscopy may arise from femtosecond laser spectral bandwidth"

### Supplemental Figure 1

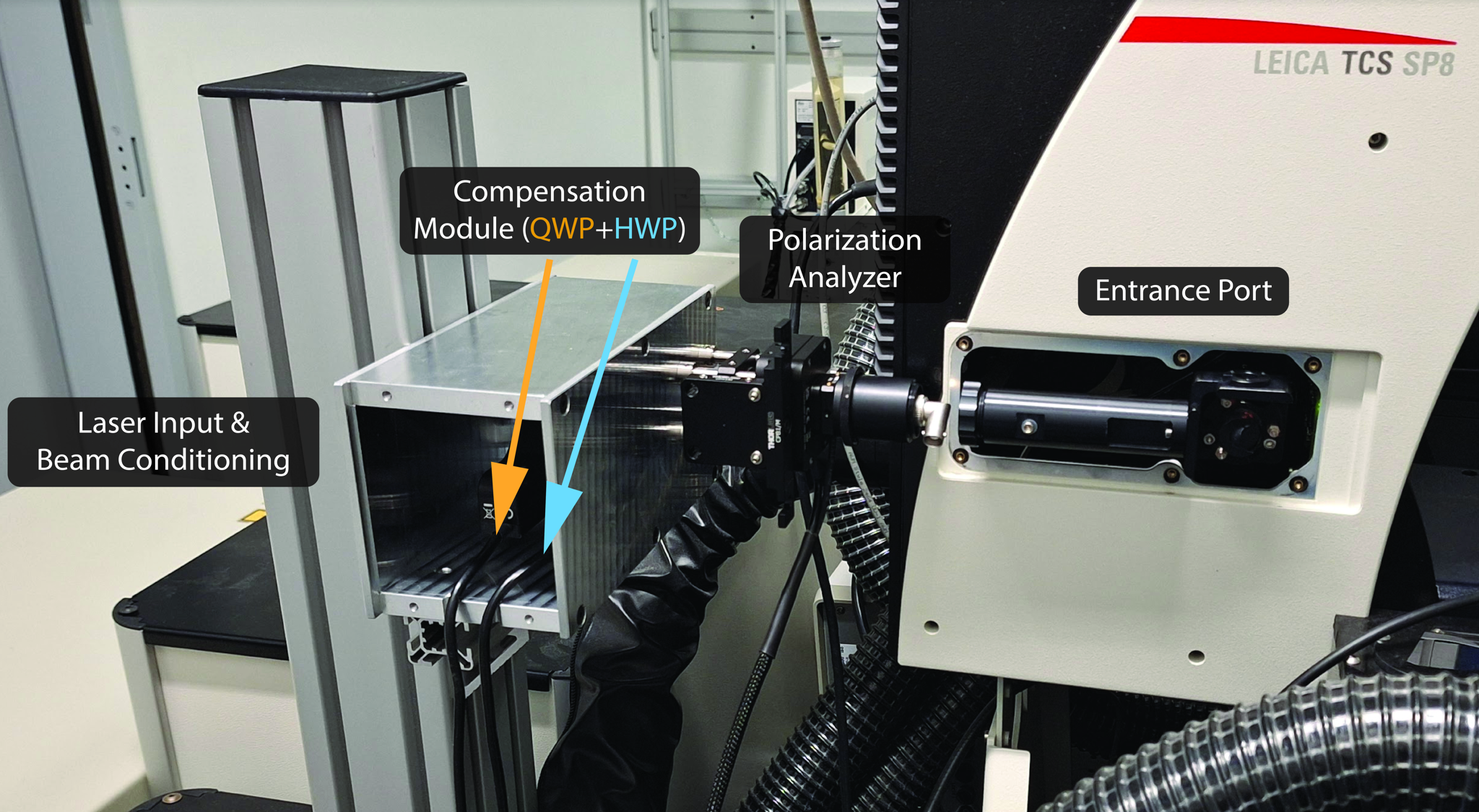

### Supplemental Figure 2

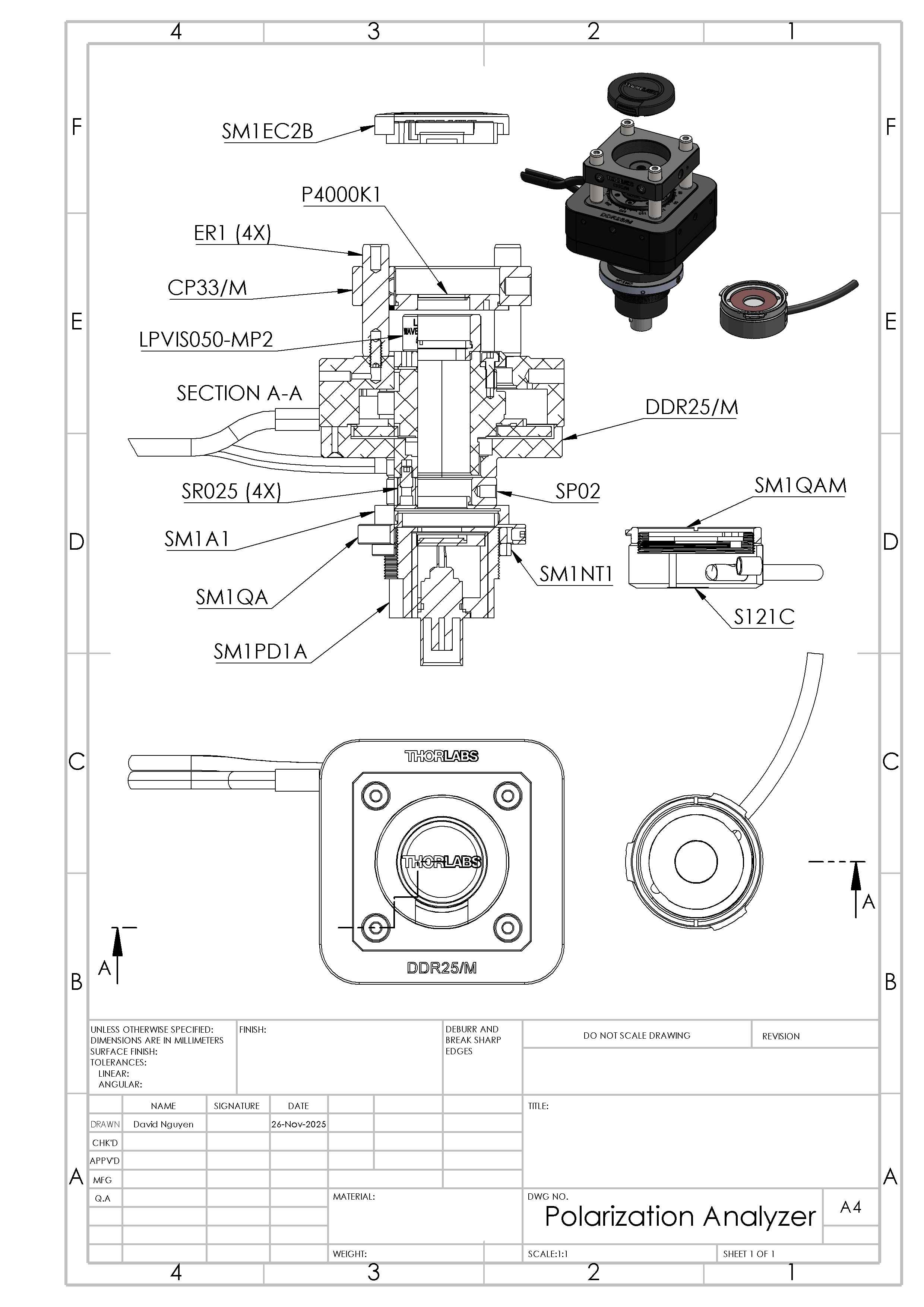

### Supplemental Figure 3

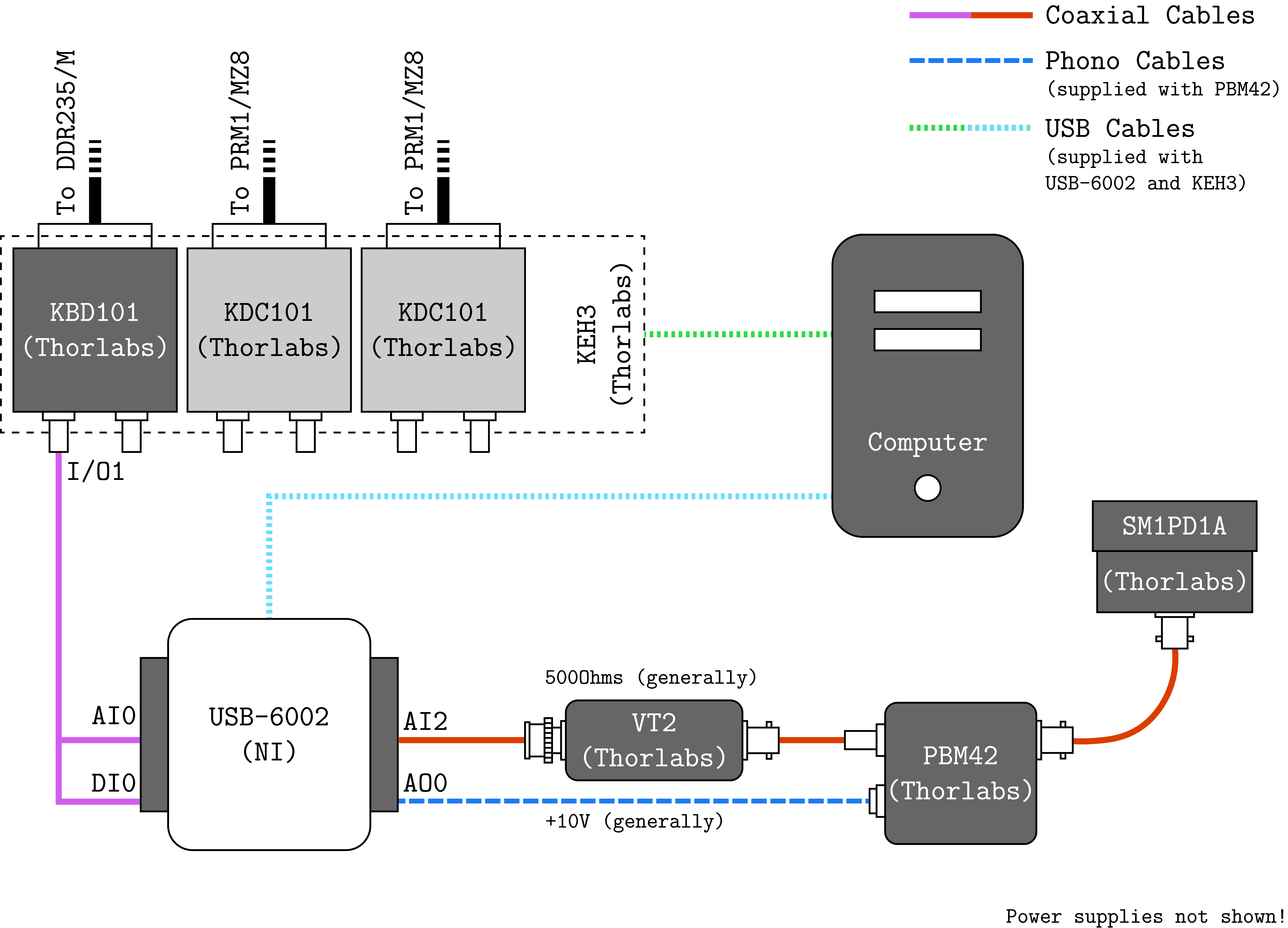

### Supplemental Figure 4

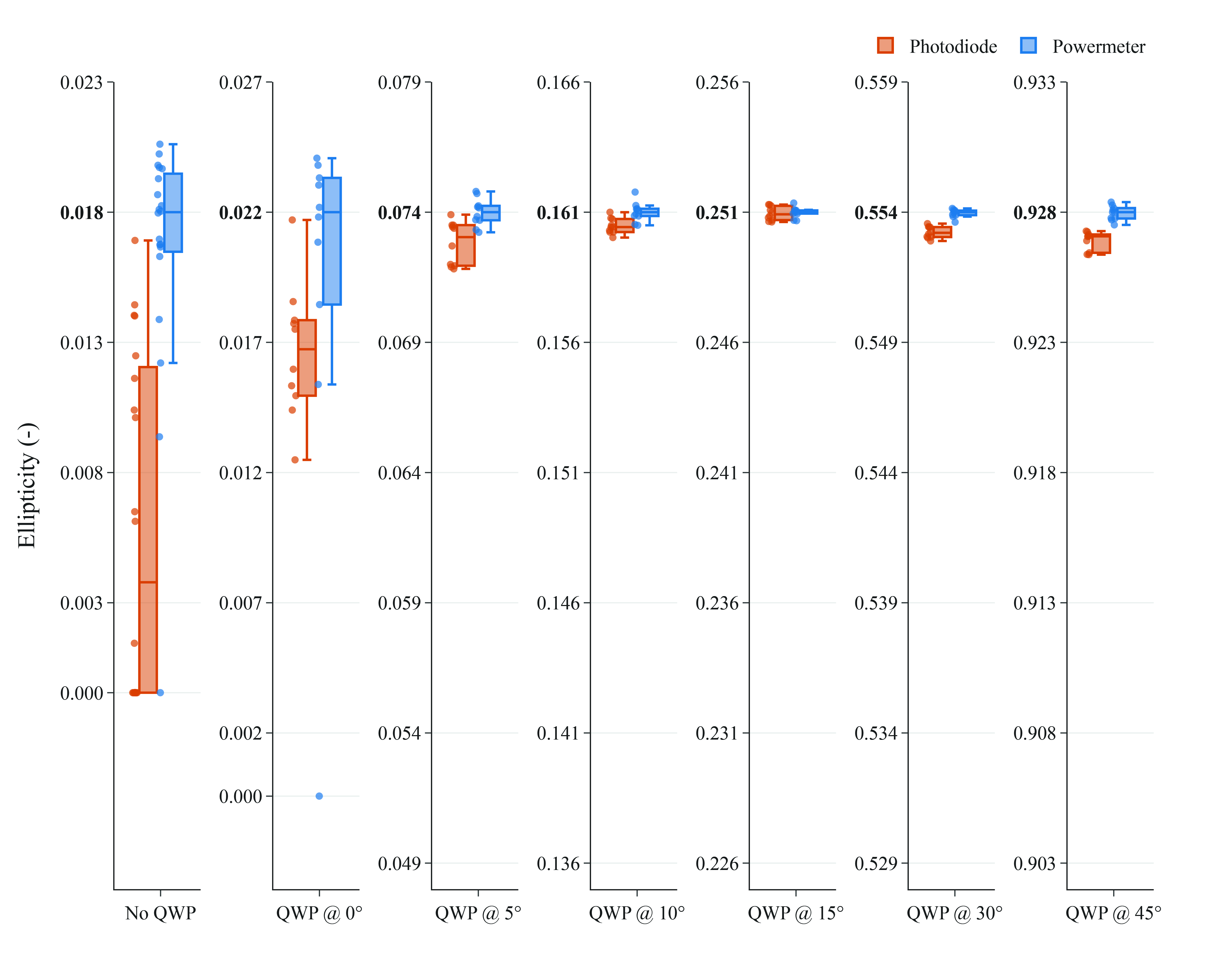

### Supplemental Figure 5

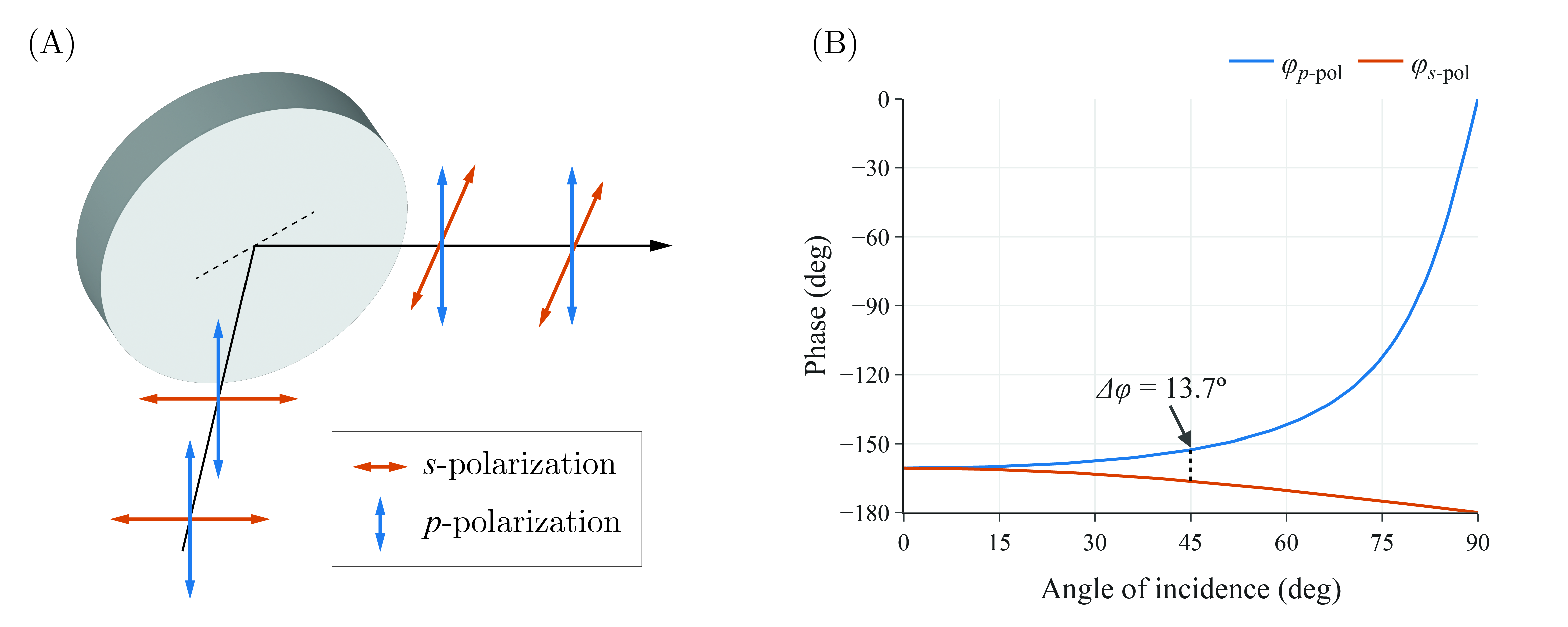
